# BAF chromatin remodeling complex subunit diversity promotes temporally distinct gene expression programs in cardiogenesis

**DOI:** 10.1101/166983

**Authors:** Swetansu K. Hota, Jeffrey R. Johnson, Erik Verschueren, Reuben Thomas, Aaron M. Blotnick, Yiwen Zhu, Xin Sun, Len A. Pennacchio, Nevan J. Krogan, Benoit G. Bruneau

## Abstract

Chromatin remodeling complexes instruct cellular differentiation and lineage specific transcription. The BRG1/BRM associated factor (BAF) complexes are important for several aspects of differentiation. We show that the catalytic subunit *Brg1* has a specific role in cardiac precursors (CPs) to initiate cardiac gene expression programs and repress non-cardiac expression. Using immunoprecipitation with mass spectrometry (IP-MS), we determined the dynamic composition of BAF complexes during mammalian cardiac differentiation, and identified BAF60c (SMARCD3) and BAF170 (SMARCC2) as subunits enriched in CPs and cardiomyocytes (CM). *Baf60c* and *Baf170* co-regulate gene expression with *Brg1* in CPs, but in CMs control different gene expression programs, although still promoting a cardiac-specific gene set. BRG1, BAF60, and BAF170 all modulate chromatin accessibility, to either promote accessibility at activated genes, while closing up chromatin at repressed genes. BAF60c and BAF170 are required for proper BAF complex composition and stoichiometry, and promote BRG1 occupancy in CM. Additionally, BAF170 facilitates expulsion of BRG1-containing complexes in the transition from CP to CM. Thus, dynamic interdependent BAF complex subunit assembly modulates chromatin states and thereby directs temporal gene expression programs in cardiogenesis.

**Significance statement:** BRG1/BRM associated factors (BAF) form multi-subunit protein complexes that reorganize chromatin and regulate transcription. Specific BAF complex subunits have important roles during cell differentiation and development. We systematically identify BAF subunit composition and find temporal enrichment of subunits during cardiomyocyte differentiation. We find the catalytic subunit BRG1 has important contributions in initiating gene expression programs in cardiac progenitors along with cardiac-enriched subunits BAF60c and BAF170. Both these proteins regulated BAF subunit composition and chromatin accessibility and prevent expression of non-cardiac developmental genes during precursor to cardiomyocyte differentiation. Mechanistically, we find BAF170 destabilizes the BRG1 complex and expels BRG1 from cardiomyocyte-specific genes. Thus, our data shows synergies between diverse BAF subunits in facilitating temporal gene expression programs during cardiogenesis.

## INRODUCTION

Cell differentiation and organogenesis are regulated by the precise transcriptional output of a coordinated gene regulatory network (1). During mammalian development, gene expression programs are spatially and temporally controlled with specific sets of genes being expressed while others are poised or repressed in a developmental stage-dependent manner. It is considered that differentiation proceeds as a gradually increasing specification of cell fates (2). Delineating the factors that control crucial developmental decision points is essential to understand the control of gene regulation during differentiation and development.

Transcription factor (TF) activity regulates transcriptional output, and is intimately influenced by the underlying chromatin. Chromatin remodeling complexes are multi-subunit protein complexes that alter histone-DNA contact in nucleosomes to reorganize chromatin and regulate transcription (3–5). The BRG1/ BRM associated factor (BAF) chromatin remodeling complexes are composed of the mutually exclusive Brahma (BRM) or Brahma-related gene 1 (BRG1) ATPAses along with several other structural subunits and their isoforms, to form diverse BAF complexes that serve specific functions in widely different cell types and developmental processes (6–8). BAF complexes have a specific composition in certain cell types; for example, an embryonic stem cell complex that regulates pluripotency (9, 10), or a neural precursor and neural BAF complexes, that fine-tune neurogenesis (11, 12). Current evidence indicates that a shift in isoforms of non-essential BAF complex subunits is important for the stepwise transition from a precursor to a differentiated state in neural and muscle differentiation (11, 13–16). Mutations in genes encoding BAF complex subunits have been associated with various cancers (17), and some have also been found to be mutated in cases of congenital heart disease (CHDs) (18). In the developing heart, subunits of the BAF complex are involved in diverse aspects of cardiac development (19, 20). *Brg1* is haploinsufficient in the mouse heart, and genetically interacts with genes encoding DNA-binding transcription factors associated with CHDs, indicating a potentially general importance of BAF complexes in these common birth defects (21).

Differentiation is thought to proceed during development as a continuous but highly regulated series of milestones, which include lineage decisions and linear progression towards a terminally differentiated state. These sequential events can be modeled effectively using pluripotent cells that are subjected to well-defined differentiation protocols (22, 23). Cardiac differentiation is composed of a stereotyped set of steps, with the initial formation of cardiogenic mesoderm, subsequent specialization into multipotent cardiac precursors, and then to their differentiation into beating cardiomyocytes (24–27). The in vivo embryonic steps are well recapitulated in in vitro differentiation protocols (25, 26). It is not known which BAF complex subunits are essential for controlling temporal steps in cardiac differentiation.

Here, we define dynamic BAF complex composition during mouse cardiomyocyte differentiation. We identify the BAF subunits BAF60c and BAF170 as enriched in cardiac precursors and cardiomyocytes. BRG1 initiates cardiac gene expression programs in precursor cells, a role shared by BAF60c and BAF170, which also maintain the cardiac program to facilitate cardiomyocyte differentiation. BAF60c and BAF170 also regulate BAF complex composition, stoichiometry and chromatin accessibility. Further, we find that BAF170 destabilizes and facilitates dissociation of BRG1 complexes from their binding sites upon cardiomyocyte differentiation. These results reveal the instructive nature that specific combination of BAF subunits attain to dictate functional outcomes during lineage commitment and differentiation.

## RESULTS

### Brg1 initiates cardiac gene expression programs during cardiomyocyte differentiation

To model early heart development, we differentiated mouse embryonic stem cells (mESCs) to cardiac troponin T positive (cTnT+), beating cardiac myocytes (25, 26). To understand the role of the BAF complex ATPase BRG1 during cardiomyocyte differentiation, we conditionally deleted *Brg1* (also known as *Smarca4*) using an inducible Cre-loxP system (Fig 1A) (9, 28). Loss of *Brg1* at CP, but not CM, inhibited cardiomyocyte differentiation (Fig. 1B). RNAseq revealed that *Brg1* regulated a total of 545 genes in CP and 125 genes in CM (*p* <0.05, ± 1.5-fold). Reduced importance of *Brg1* at the cardiomyocyte stage is consistent with its reduced expression (Fig. 1C and S1A). In CPs, *Brg1* repressed 197 (36.1%) and activated 348 (63.9%) genes (Fig. 1C). BRG1 activated genes were enriched for sarcomere organization and assembly and are essential components of cardiac cell fate establishment (Fig. 1D). Of note, *Brg1* is essential for the immediate activation of these lineage-specific genes, in anticipation of the final differentiation status of the cells, but not their maintenance. Thus, *Brg1* primes the cell-type specific differentiation of cardiac precursors. In CMs, although BRG1 activated and repressed roughly equal number of genes, it did not enrich for any biological processes, consistent with in vivo data (29).

**Fig. 1.**
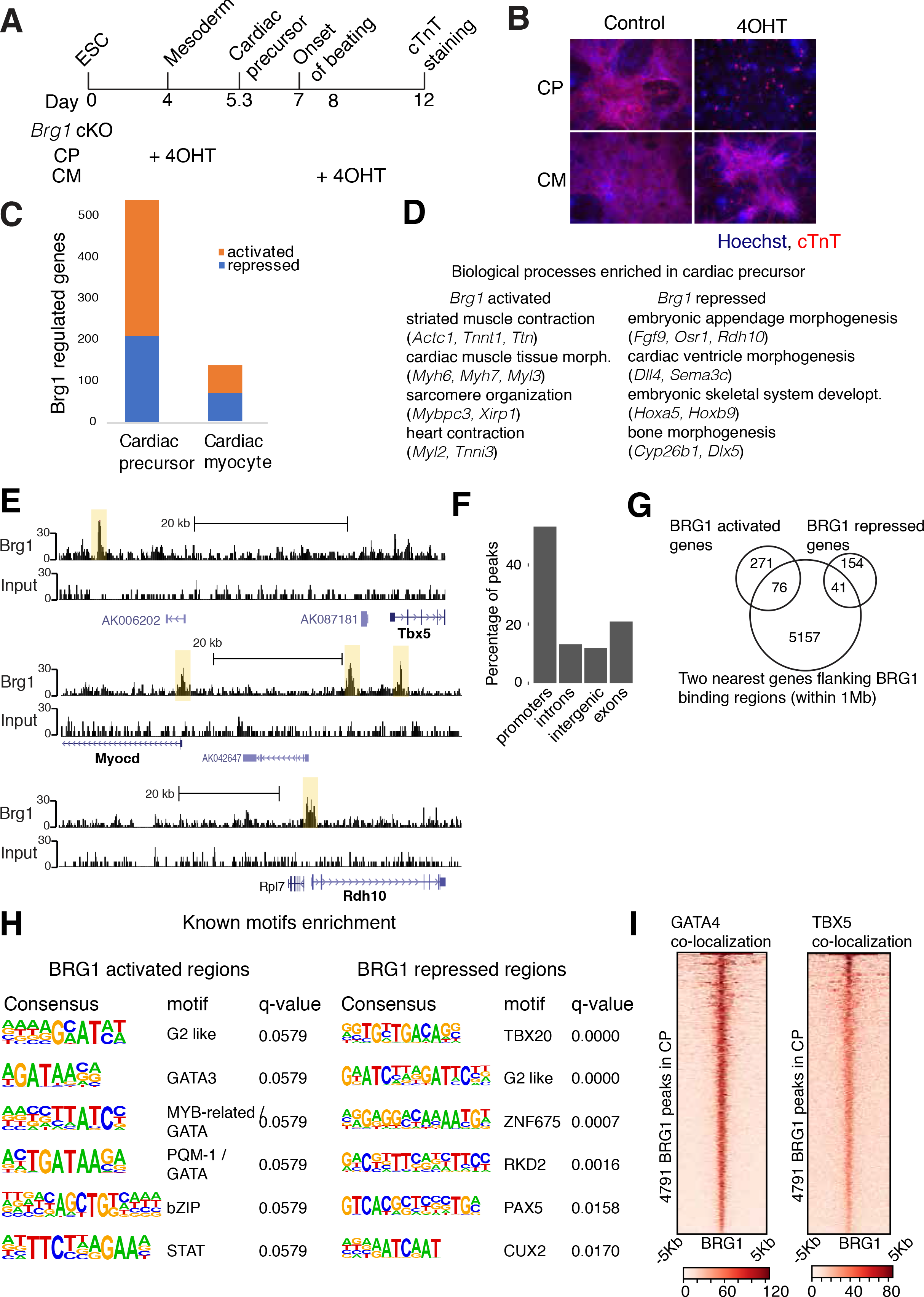
BRG1 directs cardaic gene expression. (*A*) Schematics of cardiac differentiation showing timing of 4-hydroxyl tamoxifen (4OHT) addition to conditionally delete *Brg1.* (*B*) Immunofluorescence of cTnT at D10 of differentiation. (*C*) RNAseq of BRG1 regulated genes at CP and CM. (*D*) Gene ontology (GO) biological processes enriched in BRG1 activated and repressed genes in CPs. (*E*) Input and BRG1 ChIPseq browser tracks in CPs. (*F*) Classification of BRG1 ChIPseq peaks into different genomic regions. Promoters are defined as within 1 Kb of TSS. (*G*) BRG1 activated or repressed genes overlapping with two nearest genes flanking BRG1 binding sites shown in a Venn diagram. (*H*) Motifs enriched in BRG1 binding sites associated with BRG1 activated genes and repressed genes. (*I*) GATA4 and TBX5 binding correlates well over BRG1 binding sites in cardiac precursors.

### BRG1 binding correlates with sites of cardiac transcription factor binding

To distinguish between direct vs indirect role of BRG1 in gene regulation, we performed ChIPseq using a native antibody against BRG1. In CPs, we found 4791 significant BRG1 binding regions (Figs. 1E and S1B) and could detect weaker BRG1 occupancy in CMs (Fig. S1C), presumably due to the low levels of protein at this stage. In CPs, BRG1 was bound to both transcriptional start sites (TSS) and H3K27ac marked active enhancer regions (26) (Figs. 1F and S1D) and associated with 5274 genes flanking BRG1 binding sites (within 1Mb) (Fig 1G). 21.9% of BRG1-activated and 20.9% of the BRG1-repressed genes overlapped with genes nearest to a BRG1 binding site. Putative BRG1-activated direct targets enriched for cardiac muscle development, contraction and circulatory system processes whereas BRG1-repressed direct targets enriched for embryonic limb and skeletal system development (Fig. S1E). These results show that BRG1 facilitates of cardiac gene expression programs while preventing other developmental programs including embryonic limb development.

Motif enrichment analysis using HOMER (30) revealed GATA motifs near BRG1 activated sites and T-box motifs near BRG1 repressed sites among others (Fig. 1H). Consistently, BRG1 binding sites correlated well with GATA4 and TBX5 binding sites in cardiac precursors (Fig. 1I) (31), suggesting that BRG1 interacts with cardiac TFs to regulate gene expression during cardiac lineage commitment, as predicted from gain of function experiments (32, 33). Thus, BRG1 (and BRG1 containing complexes), in collaboration with cardiac transcription factors, direct cardiac gene expression program while preventing expression of non-cardiac genes.

### BRG1 complex shows dynamic composition during cardiomyocyte differentiation

To understand the composition of BRG1-associated complexes during cardiac differentiation, we immunoprecipitated BRG1 under stringent conditions using a BRG1-3x FLAG heterozygous mouse embryonic stem (ES) cell line (34), from five different stages of cardiac differentiation: ES cells (ESC), embroid bodies (EB), mesoderm (MES), cardiac progenitor (CP) and cardiac myocytes (CM) (Fig. 2A). An untagged control cell line similarly processed in biological triplicates at each stage served as negative control. BRG1 complexes isolated from each of these stages showed overall protein profiles similar previously reported complexes (Fig. 2A) (10). We performed mass-spectrometry in biological triplicates and technical duplicates and compared peptide intensities after normalizing against untagged control, the bait BRG1 protein, and across stages of differentiation, and identified the composition of the BRG1 complexes (Fig. 2B and Table S1). ESC-derived BRG1 complexes were enriched for BRD9, GLTSCR1l, BCL7b&c, BAF45b, BAF155 and BAF60a, consistent with previous reports (10, 35). We identified proteins enriched at mesoderm (PDE4D, CRABP2, ARID1B), CP (BAF60b, POLYBROMO-1, ARID2, BAF47, BCL7a, BRD7, and BAF45a) and CM stages (WDR5, BAF170, BAF60c, BAF57, SS18l1, BAF45c and CC2D1B). New BRG1 interacting factors were identified, including CRABP2, KPNA2, PDE4D, CSE1L, CC2D1B, and intriguingly, WDR5. WDR5 is well known to be part of the MLL complex, and has been implicated in human CHD (36). Glycerol gradient experiments showed co-sedimentation of BRG1-associated subunits in ES cells and cardiomyocytes confirming intact nature of the isolated complexes, and the inclusion of WDR5 (Figs. 2C and 2D). Functional assessment of BRG1 complexes by in vitro nucleosome repositioning or ATPase activity assays indicated that stage-specific complexes had similar activities (Figs. S2A-C). Thus, BRG1 complexes have dynamic subunit composition that gradually changes during cardiac differentiation and enrich for specific subunits in cardiac lineages.

**Fig. 2.**
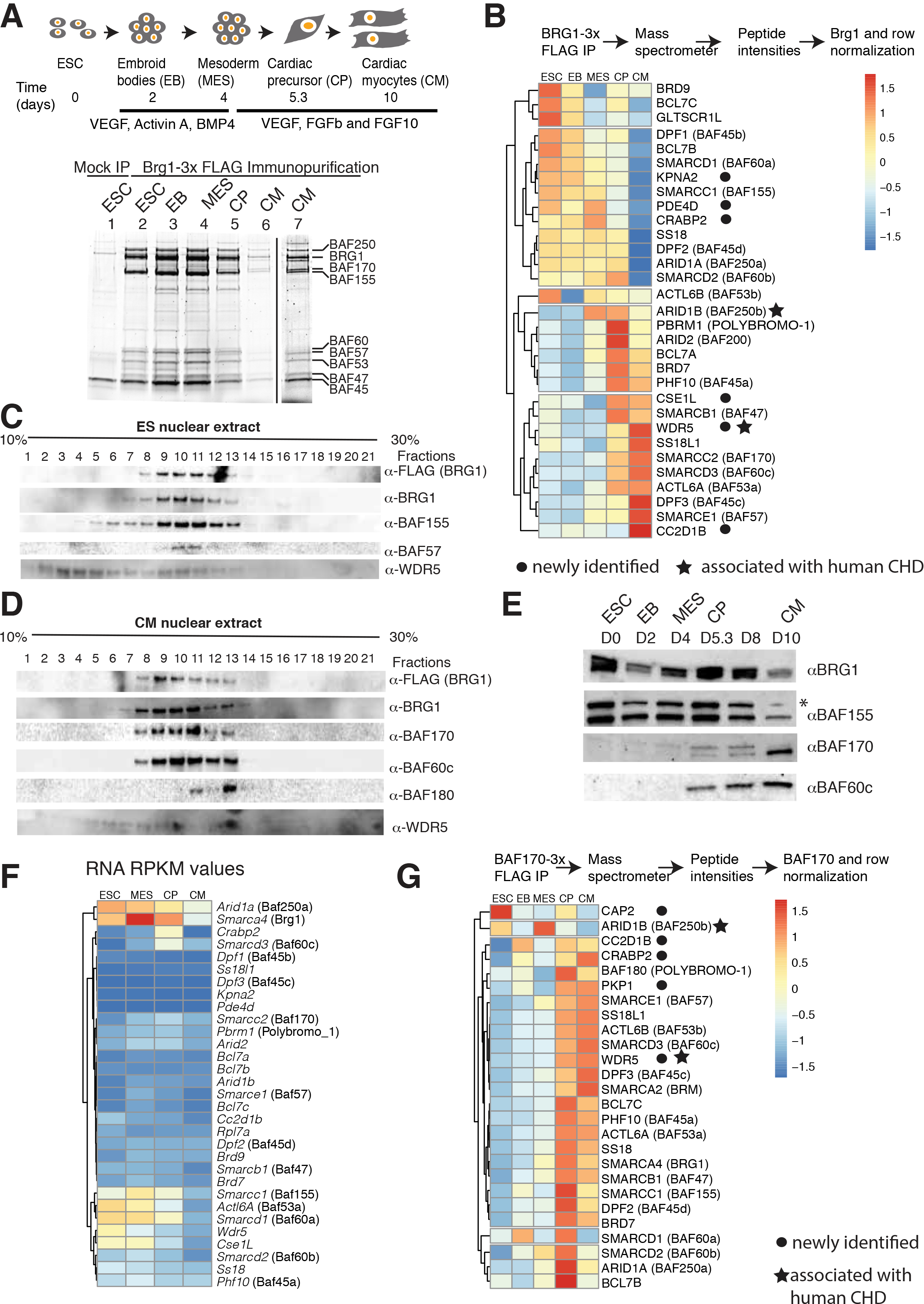
BRG1-containing complexes dynamically change subunit composition during cardiac differentiation (*A*) Schematic representation of cardiac differentiation and isolation of BRG1 complexes by anti-FLAG immunoprecipitation. SDS-PAGE showing BRG1 complexes from an untagged (lane 1) and a Brg1-3x FLAG tagged mESC line differentiated to cardiomyocytes at different stages (lane 2-6). BRG1 complex isolated at CM (lane 6) is shown with increased contrast (lane 7). Protein complexes were analyzed in a 10% SDS-PAGE and stained with SyproRuby protein gel stain. (*B*) Mass spectrometric analysis of BRG1 complexes. The peptide intensities are normalized to mock untagged intensities, BRG1 intensities and further normalized across row. Color blue and red indicates depletion and enrichment respectively while yellow represent unchanged. Nuclear extracts of ESC (*C*) or CM (*D*) resolved in a 10-30% glycerol gradient and BRG1-associated subunits detected by western blot. (*E*) Nuclear extracts from different stages of cardiac differentiation detected BAF/PBAF subunits by western blot. (*F*) RNAseq expression profile (RPKM) of BAF/PBAF subunits during cardiac differentiation. (*G*) Peptide intensities of anti-FLAG BAF170-3x FLAG protein after IP-MS, normalized to mock, BAF170 proteins and across stages of differentiation. Color scheme same as panel B.

These results suggest that BRG1 complex changes its composition during cardiac differentiation and BAF subunits switch from one isoform to other (for example, BAF60a in ES is replaced by BAF60c in CP/CM BAF complexes) or to a different protein (BAF155 in ES cells to BAF170 in cardiac cells). Switch from BAF155 in ES BAF to more abundant BAF170 in CP/CM and appearance of BAF60c only in cardiac cell lineages during differentiation were consistent with their expression pattern (Fig. 2E). However, most subunits were not developmentally regulated, and thus the assembly reflects developmental stage-specific inclusion of these subunits (Fig 2F). For example, WDR5 is expressed mostly in ESC, while it associates with BAF complexes only in CMs.

To further understand the composition of cardiac enriched complexes, we immunoprecipitated endogenous BAF170-3xFLAG, at five different stages of cardiac differentiation in biological triplicates, again with an untagged control line for each stage, and analyzed immunoprecipitated proteins by mass spectrometry. Most of the proteins enriched at the CP and CM stages using BRG1 as bait were also enriched using BAF170 as bait, with certain exceptions (Fig. 2G). For example, ARID1b is enriched in the BRG1 IP-MS at the MES-CM stages, while in the BAF170 IP-MS it is depleted. The opposite dynamic enrichment is seen for CRABP2. WDR5 was also present in the complexes isolated by BAF170-IP. In addition, we detected the alternate ATPase BRM in the BAF170-FLAG purification, indicating that BAF170 functions within separate BRG1 and BRM complexes. These results suggest that dynamic subunit composition and subunit switch are important aspects of BAF complexes during cardiac differentiation.

### BAF170 and BAF60c facilitate cardiomyocyte differentiation

We investigated the functional roles of cardiac BAF enriched subunits BAF60c and BAF170 by deleting them in ES cells using TALEN or CRISPR strategies (Fig 3A). Both BAF60c KO and BAF170 KO cells underwent cardiac differentiation as observed by beating cardiac myocytes and immunostaining of cardiac Troponin T (Fig. 3B). Cells lacking BAF170 however had a delay in onset of beating (Fig 3C) and both BAF60 and BAF170 KO cells displayed aberrations in beating (Fig 3D), indicative of abnormal cardiac differentiation. To understand how these subunits regulate gene expression we collected cells at CP and CM stages and performed RNAseq. In CPs, BAF60c regulated a total of 474 genes (p-value <0.05, ± 1.5-fold) of which 45% were activated and 55% were repressed, while a total of 382 genes are deregulated in BAF170 KO, of which roughly equal number of genes were up or down regulated (Fig S3A). Gene ontology analyses revealed that BRG1 and BAF60c share common functions in activating genes involved in cardiac and muscle tissue development and contraction while repressing skeletal system and limb morphogenesis (Fig S3C). Genes regulated (largely repressed) by BAF170 are mostly involved in skeletal system morphogenesis and pattern specification (Fig. S3C). However, in CMs, BAF60C regulated a large number of genes in cardiomyocytes (2646), repressing 72% of genes and activating 28% of these genes (Fig S3B). Similarly, BAF170 also repressed a large percentage (63%) of genes in CMs (Fig. S3B). These results indicate a largely repressive function of these cardiac enriched subunits in gene regulation.

**Fig. 3.**
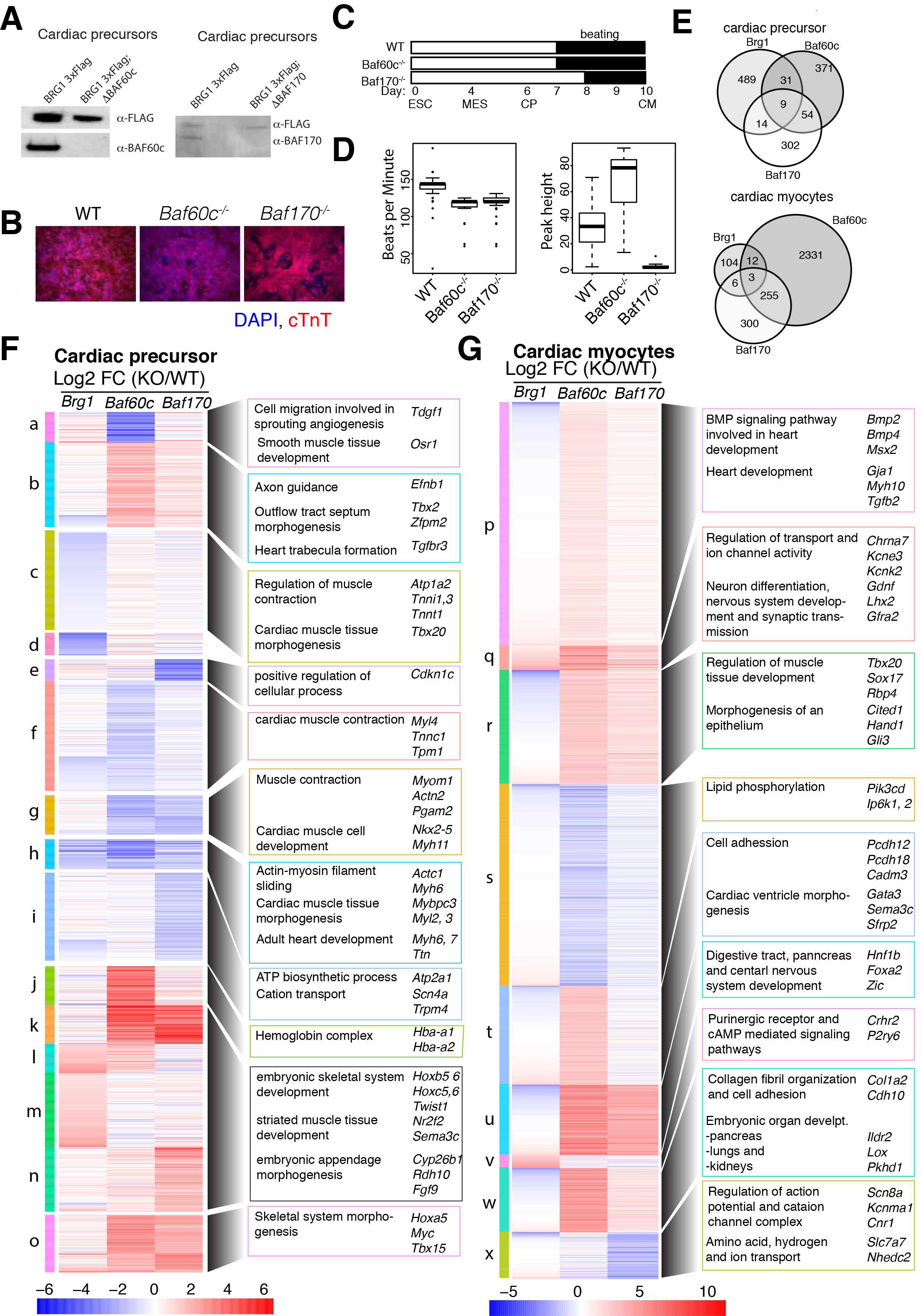
BAF60c and BAF170 regulate both common and distinct gene expression programs compared to BRG1. (*A*) Loss of BAF60c and BAF170 proteins in respective KO cells detected by western bot in cardiac precursor cells. BRG1 levels were shown as loading controls. (*B*) Immunofluorescence of cardiac troponin T protein in WT, BAF60cKO and BAF170 KO cells in CMs. (*C*) Schematics showing onset of beating in differentiating cardiomyocytes. (*D*) Assessments of beating properties of WT, BAF60c KO and BAF170 KO cardiomyocytes by measuring number of beats per minute and amplitude of beating represented by peak heights. (*E*) Venn diagram shows an overlap of significantly affected genes (1.5 fold, p-value<0.05) in absence of *Brg1*, *Baf60c* and *Baf170* at CP and CM. Heat map showing comparison of genes significantly deregulates in any of the three genotypes over their respective WT counterpart at CP (*F*) and CM(*G*). Clusters are shown in vertical colored bars to the left and representative genes involved in various biological processes are shown to right of heatmap.

Overall comparison of the gene expression changes in CPs and CMs lacking BRG1, BAF60c, or BAF170, show that all three have shared functions in CP gene expression, but that in CPs and CMs BAF60c and BAF170 repress a cohort of genes independent of BRG1 (Figs. 3E-G). This clearly indicates divergent role of these two subunits from BRG1, suggesting their association in a different BAF complex or potential independent function outside of BAF complexes.

### BAF subunits modulate temporal chromatin accessibility and facilitate cardiac gene expression programs

To understand the function of BRG1, BAF60c and BAF170 in gene expression regulation, we used ATACseq (37) to examine chromatin accessibility genome-wide in CPs and CMs. We compared differential chromatin accessibility profiles in *Brg1* conditional, *Baf60c*^−/−^ and *Baf170*^−/−^ KOs in CP and CM cells. In CPs, BRG1 maintains chromatin accessibility near genes involved in cardiovascular development, cardiac tissue morphogenesis and regulation of cell differentiation (Fig 4A, clusters *a*, *g* and *i*) and prevents chromatin accessibility near genes involved in transcriptional regulation and chromatin organization (Fig 4A, cluster *e*). In the absence of BAF60c, accessibility was increased at genes are involved in non-cardiac cell differentiation and cell migration. In contrast, absence of either BAF60c or BAF170 accessibility was reduced near genes involved in cardiovascular development (*Gata4, Tbx5, Myocd, Myh7*) (Fig 4G), calcium handing (*Ryr2*) and muscle contraction (*Scn5a, Scn10a, Kcnq1*). Loss of BAF60c and BAF170 increased chromatin accessibility near genes involved with embryonic limb and skeletal muscle development (*Myf5*, *Myf6*, *Tbx2*) (Fig 4E) and early embryo development (*Cer1, Dkk1, Fgf9 and Gsc*).

**Fig. 4.**
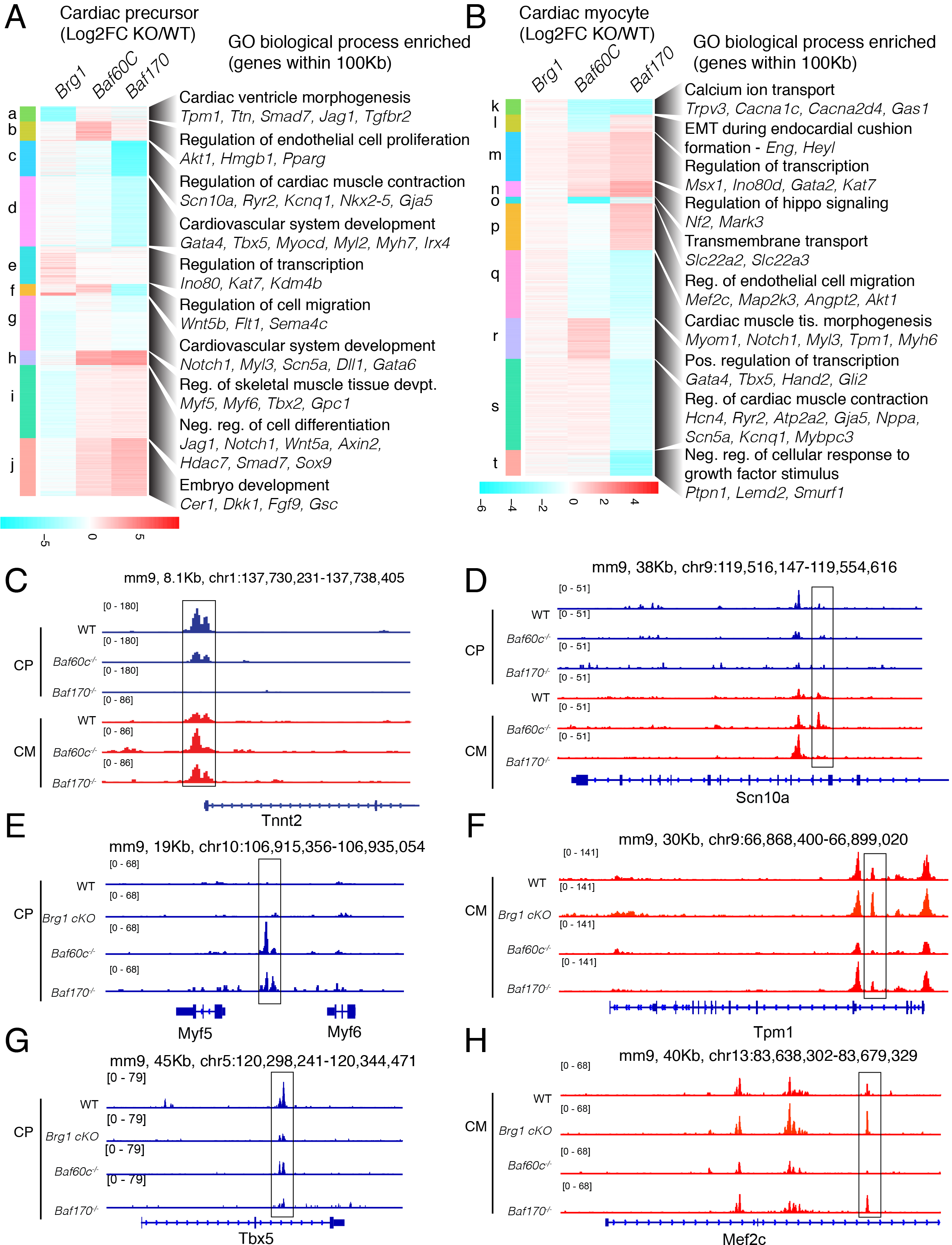
BRG1, BAF60c and BAF170 regulate temporal chromatin accessibility (A,B) Heat map of significantly affected ATAC-seq peaks (FDR <0.05, 2-fold change) in BRG1 conditional, BAF60c KO and BAF170 KO over their respective WT counterparts in CPs (*A*) or CM (*B*). Enriched biological processes of two nearest genes within 100kb of the ATACseq peaks were analyzed using GREAT and shown to the right with representative genes. Browser tracks showing delayed opening of *Tnnt2* promoter chromatin (*C*) and loss of *Scn10a* enhancer in absence of BAF170 (*D*). Chromatin opening and closing near *Myf5* (*E*) and in *Tbx5* (G) in absence of BAF60c and BAF170 in CPs and chromatin closing in *Tpm1* (F) and *Mef2c* (H) in absence of BAF60c in CMs are shown.

In CMs, loss of BRG1 did not change chromatin structure, consistent with gene expression (Figs 4B and 1C), while loss of BAF60c and BAF170 increased chromatin accessibility near genes involved in chromatin and transcription regulation (*Ino80d, Kat7*) and hematopoietic differentiation (*Gata2*). BAF60c alone is implicated in promoting chromatin accessibility near cardiac precursor genes (*Gata4, Tbx5, Hand2*), while BAF60c and BAF170 are important in chromatin accessibility near genes involved in cardiac function (*Myom, Tpm1, Myh6 and Myl3*) (Fig 4F) and calcium ion transport (*Cacna1c, Cacna2d4*). Uniquely they also regulated chromatin near signaling and cell migration genes (Fig 4H). Further, in absence of BAF170 we observed delayed accessibility of the *Tnnt2* promoter (Fig 4C) consistent with delayed onset of beating (Fig 3C), and impaired accessibility at an enhancer in the *Scn10a* gene (Fig 4D). This *Scn10a* region is a known TBX5-dependent enhancer of *Scn5a*, and was identified as containing a GWAS SNP associated with altered electrophysiology in humans (38, 39). These data provide evidence of the unique and shared functions that individual BAF subunits exert to regulate chromatin structure to facilitate a cardiac gene expression program while maintaining repression of non-cardiac developmental genes.

### Both BAF170 and BAF60c regulate BAF complex stoichiometry

BAF60c and BAF170 have critical roles in regulation of chromatin structure and transcription. It is not clear whether mammalian BAF complex composition is reliant on the presence of specific subunits. To understand the nature of BAF complexes formed in absence of these cardiac enriched subunits, we Immunoprecipitated BRG1-3xFLAG complexes from ES cell lines lacking BAF60c or BAF170 at CP and CM stages. SDS PAGE revealed increased stability of BRG1-containing complexes in BAF170 KO CMs (Fig. 5A). During the CP to CM transition, BRG1 complex abundance is reduced in WT (Fig 2A, compare lane 5 to lane 6) and BAF60c KO (Fig. 5A, compare lanes 2 - 4 to lanes 5 & 7) cells. In BAF170 KO the abundance of BAF complexes remained unchanged (Fig. 5A, compare lanes 8 & 9 to lanes 10 & 11), indicating a greater stability of BRG1 complexes in BAF170 KO cells. MS analyses of these complexes revealed significant differences in subunit composition in the absence of BAF60c or BAF170 (Fig. 5B,5C). In CPs, BRG1 complexes lacking BAF60c had reduced association of BAF53a, BRD7, BAF45a and BAF45c and were enriched for BAF45d, BAF47, SS18, BAF155 and BAF60a (Fig. 5B and S5A). In CMs, we observed increase association of many subunits with BRG1 in absence of BAF60c (Figs. 5B, S4A and S4B). Similarly, BAF170 loss had reduced association of BCL7a, BAF45d, BAF60c, BAF47 and BAF53a, and increased association of BAF45c, CREST and BAF180 with BRG1 in CPs. In CMs, loss of BAF170 reduced association with BAF60c and increased association with BAF155 and BAF60b (Figs. 5C, S4A and S4B). The altered complexes formed are not due to changes in the transcriptional level of BAF subunits (Fig. S4C).

**Fig. 5.**
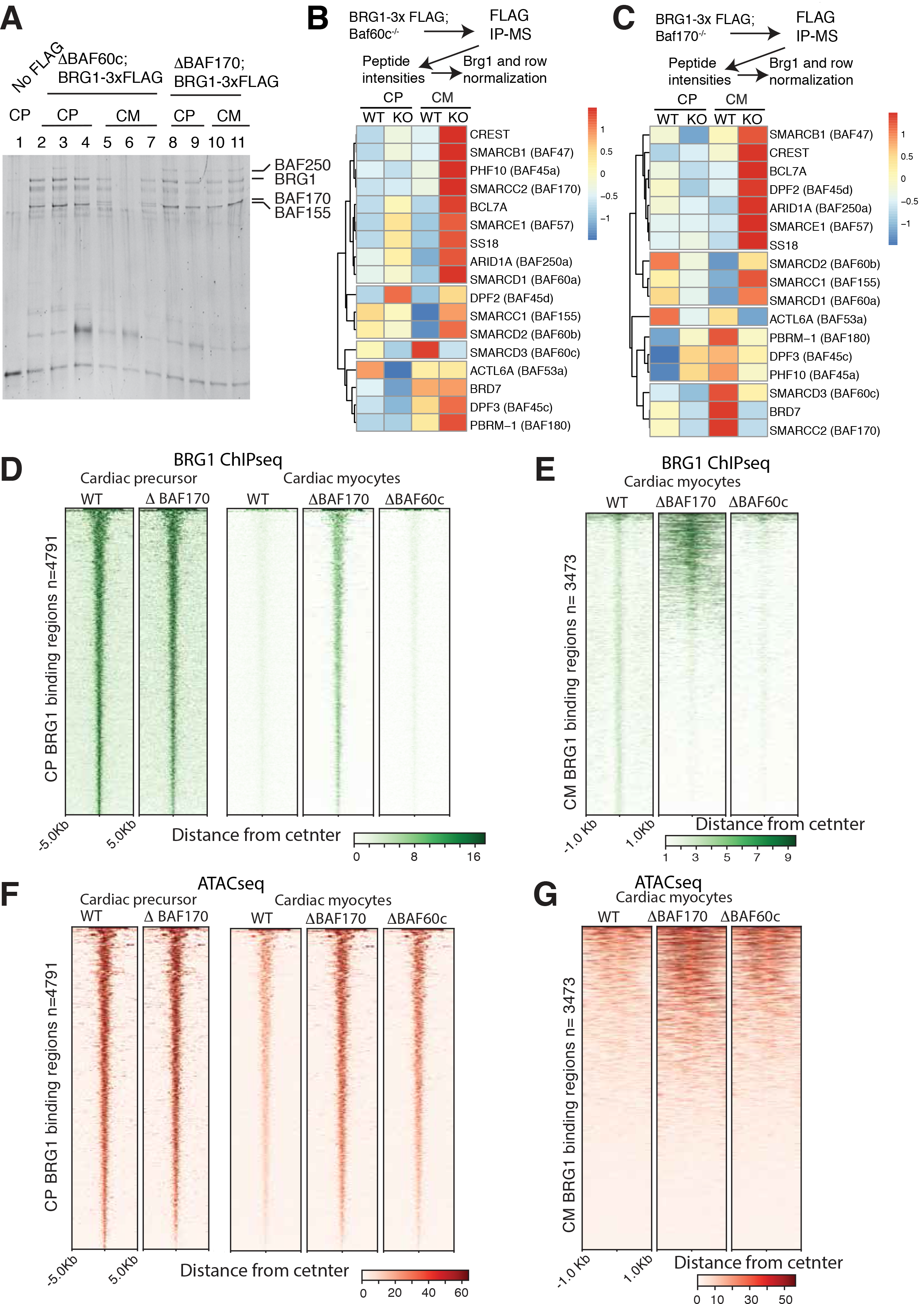
Cardiac enriched BAF subunits regulate BRG1 complex stoichiometry, stability and genome binding. (*A*) Immunoprecipitated and FLAG eluted BRG1-containing complex isolated from WT or cells lacking of BAF60c or BAF170 in CPs and CMs and analyzed on a 10% SDS-PAGE. (*B*) Peptide intensities of FLAG IP-MS in absence of BAF60c (*B*) or BAF170 (*C*) in CP and CM after normalization to mock control, BRG1 levels and across stages of differentiation. (*D*) ChIP-Seq showing BRG1 binding over 4791 peaks in WT and BAF170 KO in CPs (left panel). BRG1 binding over these same 4791 CP sites in WT, BAF170 KO or BAF60c KO in CMs (right panel). (*E*) BRG1 ChIP over 3473 peaks in WT, BAF170KO and BAF60c KO in CMs. (G) ATACseq signal over 4791 BRG1 binding sites in CP and CM for the indicated genotypes. (H) ATACseq signal over 3473 CM BRG1 binding sites over indicated genotypes in CM.

These results suggest that a fine balance exists in the composition of the BAF complex and perturbation of one subunit extends to the stoichiometry of association of other subunits in the complex. It further indicates that different subunits or isoforms substitute for the absence of one or more subunits. For example, both BAF60a and BAF60b substitute for the lack of BAF60c, and depletion of BAF45c is balanced by enrichment of BAF45a (Fig. S4B). These results emphasize that subunit switching and substitution in BAF complex composition could be an important mechanism in cardiac lineage specification.

### BAF170 facilitates temporal BRG1 dissociation from the genome

We explored the possibility that BAF60c or BAF170 help direct the genomic localization of BRG1. In CPs, BRG1 binds to a set of 4791 sites (Fig. S1B) and these are largely unaffected in absence of BAF170 (Fig. 5D). BRG1 binding at these site is severely reduced in cardiomyocytes in WT and cells lacking BAF60c, presumably due to the reduced predominance of a BRG1-containing complex. However, in the absence of BAF170, BRG1 binding at a large subset of these sites is retained in cardiomyocytes (Fig. 5D), consistent with the stabilization of BRG1-containing complexes. These chromatin regions are accessible in CPs, and normally subsequently inaccessible in CMs, but remain accessible in absence of BAF170, consistent with retained BRG1 complexes remodeling these regions (Fig. 5F). In CMs, BRG1 weakly bound to 3473 sites, and lack of BAF170 increased BRG1 binding to some of these (Fig 5E), while the absence of BAF60c reduced the binding BRG1. These binding dynamics correlated with chromatin accessibility (Fig 5G). This suggests that recruitment of BRG1-inclusive complexes is regulated by cardiac specific subunits, and that concomitant dynamic expulsion of the complex is also highly regulated by similar processes.

## DISCUSSION

BRG1 containing complexes respond to and modulate lineage decisions and differentiation in cardiac development by incorporating specific subunits at discrete stages. These specialized BAF complexes regulate distinct gene expression programs to drive cardiogenesis while simultaneously repressing alternate non-cardiac developmental programs. Our results suggest that BAF complexes use multiple interdependent mechanisms including switching subunits or isoforms, regulating subunit stoichiometry, altering BRG1 recruitment, and expulsion strategies, to modulate dynamics of cardiac differentiation. This unanticipated complexity of chromatin complex regulation emerges as a critical determinant of differentiation.

Subunit composition switching of chromatin remodeling complexes have been reported during neural development (11, 40, 41) and skeletal myogenesis (15, 16). For example, during myogenic differentiation, BAF60c expression is facilitated over the alternative isoforms BAF60a and BAF60b, which are post-transcriptionally repressed by myogenic microRNAs. However, the systematic IP-MS-based discovery of BAF/PBAF complex subunits during the time course of a well-regulated differentiation process has to date not been achieved. Importantly, we show that BRG1-associated subunits at different stages of cardiac differentiation form specific complexes to activate or repress specific transcriptional programs. In addition, we determine that the presence of specific BAF subunits greatly dictates the overall composition, and therefore function, of the complex.

The continued roles played by BAF60c and BAF170 are intriguing considering the reduced role of BRG1 in later stages of cardiac differentiation. The incorporation of these subunits within a BRM-containing complex may explain this, although the apparent absence of a developmental or postnatal cardiac phenotype in BRM null mice might indicate minimal importance of these complexes (42). It has been suggested that BAF60c may function independently from the BAF complex (43, 44), although there is no evidence for this in cardiac cells. It is also possible that the altered BAF complex assembly in the absence of BAF60c or BAF170 creates a set of complexes with anomalous function, thus leading to aberrant gene expression.

BAF60c and BAF170 are essential for BRG1 complex composition and stoichiometry. Altered association or dissociation of several subunits in the absence of BAF60c or BAF170 indicates the presence of sub-modules within BAF complex mediated by these subunits. Our observations are consistent with reports in yeast SWI/SNF complexes where absence of specific subunits form sub-modules or altered SWI/SNF complex composition (45–47). Thus, phenotypic observations in absence of a particular subunit may be the result of disruption of a module containing many subunits. Conflicting evidence regarding this mechanism has been shown for other BAF complex subunits in mammalian cells (48, 49). We interpret our results as indicating that a stable complex with broadly altered composition exists in the absence of BAF60c or BAF170.

BRG1 binding to genomic sites in CP and putative BRG1 direct targets have critical roles in promotion of cardiac gene expression programs while repressing non-cardiac fate. BRG1 binding is greatly reduced in CMs indicating diminished BRG1 function. Our finding that BAF170 largely helps BRG1 dissociation from the genome indicates a potential mechanism of BAF170 mediated gene regulation during differentiation. How BAF170 regulates genomic expulsion of BRG1 is currently not understood, but association of BAF170 with BRM containing complexes in CM might evict BRG1 from BAF complexes as both BRG1 and BRM are mutually exclusive. Further, concomitant increase in BAF170 and decrease of BAF155 association with BRG1 containing complexes during cardiac differentiation might make BRG1 prone to post-transcriptional modification and proteasome mediated degradation, as BAF155 is known to stabilize BRG1 containing complexes (50). Consistently, we observe BRG1 complex stabilization in absence of BAF170 in CMs.

In conclusion, our study identifies morphing combinations of very specific BAF subunits that form and change subunit compositions during cardiac differentiation, and drive stage-specific cardiac gene expression programs. These results are consistent with the known in vivo roles of BAF complex subunits (20, 21, 29). Genetic dissection of individual subunit contribution clearly unveils a profound and specific function of individual BAF complex subunits. How BAF chromatin remodelers with different subunit composition provide specificity to gene expression programs will be the focus of future studies.

## Author Contribution

Project design and direction: B.G.B. and S.K.H. ES cell engineering, in vitro differentiation, complex isolation, biochemistry, gene expression analysis, ATACseq, data analysis: S.K.H. Bioinformatics: R.T. and S.K.H., ES cell culture and cardiomyocyte differentiation: A.M.B. Mass spectrometry and analysis: J.R.J, E.V., under direction of N.J.K. ES cell strain construction: Y.Z. under direction of L.A.P. and X.S. Manuscript writing: S.K.H and B.G.B with contribution from all authors.

## Acknowledgements

We thank Katherine Pollard for advice and guidance on computational aspects of this project. We thank A. Williams and S. Thomas (Gladstone Bioinformatics Core) for data analysis, L. Ta and J. McGuire (Gladstone Genomics Core) for RNAseq library preparation, the UCSF Center for Applied Technologies for sequencing, B. Bartholomew, J. Persinger, G. Narlikar and C. Zhou for advice and help with nucleosome remodeling assays, and G. Howard for editing. RNAseq (258R1 and 420R), ATACseq (466R) and ChIPseq (451R) data are publicly available at https://b2b.hci.utah.edu/gnomex/ and GEO (). This work was supported by grants from the NIH/NHLBI (R01HL085860, P01HL089707, Bench to Bassinet Program UM1HL098179), the California Institutes of Regenerative Medicine (RN2-00903), and the Lawrence J. and Florence A. DeGeorge Charitable Trust/American Heart Association Established Investigator Award (B.G.B); and postdoctoral fellowships from the American Heart Association (13POST17290043) and Tobacco Related Disease Research Program (22FT-0079) to S.K.H. and NIH training grant (2T32-HL007731 26) to S.K.H. L.A.P. was supported by NHLBI grant R24HL123879, and NHGRI grants R01HG003988, and UM1HG009421, and research was conducted at the E.O. Lawrence Berkeley National Laboratory was performed under Department of Energy Contract DE-AC02-05CH11231, University of California. This work was also supported by an NIH/NCRR grant (C06 RR018928) to the J. David Gladstone Institutes and by The Younger Family Fund (B.G.B.).

## Materials and Methods

### Cardiomyocyte differentiation

Mouse embryonic stem cells (ESCs) were cultured in feeder free condition in serum and leukemia inhibitoty factor containing medium. Cardiomyocyte were differentiation as described before(25, 26). *Brg1* was deleted in presence of 200nM 4-hydroxytamoxifen for 48hrs with control cell treated similarly with tetrahydrofuran(52).

### Knockout cell line generation

BAF60c was inactivated by targeting exon 2 of *Baf60c* gene with two pairs of TALENs following Sanjana et. al (53) except a CAGGS promoter replacing the CMV promoter (54). The TALEN’s targets are as follows: GCCCCCTAAGCCCTCTCCAGAGAACATCCAAGCTA GAATGACTTGGTCGCTGCTAC and CCCGCCCCTCTCCAAGACCCTGGGTTGGTA ACCCTGCGCTGAGCGATGAGTGGGAG.

BAF170 was targeted using CRISPR/Cas9 with sgRNA targeting Exon 2 of *Baf170* gene following the protocol in Congo et al.(55). sgRNA were cloned to Bbs1 digested pX330 vector by annealing the following primers 5’ caccg CGCACCGCTTACTAAACTGC 3’ and 5’ aaac GCAGTTTAGTAAGCGGTGCG c **3’**(lowercase to create Bbs1 digestion site). Targeting vector was constructed by cloning 458 and 459bp of Baf170 DNA upstream and downstream of midpoint of sgRNA target site into Kpn1-Xho1 and BamH1-Not1 sites of pFPF (derivative of Addgene plasmid #22678, Neomycin is replaced with Puromycin cassette). 2.5ug of each of TALEN Baf60c plasmids or 2.5ug sgRNA plasmid and 20ug of BAF170 targeting constructs were used for transfection. Single clones were selected, grown, PCR genotyped and DNA sequenced.

### Nuclear extract preparation, western blot and anti-FLAG immunoaffinity purification

Nuclear extract were prepared using protocol describe before (56). Western blotting was performed using standard techniques using PVDF membranes. Primary antibodies used were anti-Brg1 (Abcam, ab110641, 1:1000), anti-FLAG (Sigma, F1804, 1:1000) and anti-BAF155 (Bethyl, A301-021A, 1:1000), anti-BAF170 (Bethyl, 1:1000, A301-39A), anti-BAF60c (Cell Signaling Technologies, 62265, 1:0000). Secondary antibodies used are Donkey anti-rabbit IRDye 800cw (Licor, 926-32213, 1:10,000), Donkey anti-mouse IRDye 800cw (Licor, 925-32212, 1:10,000), Donkey anti-goat IRDye 680cw (Licor, 925-68074-1:10,000).

For immunoaffinity purification of BAF complexes, nuclear extracts made from 10^8^ cells were incubated with 50ul of anti-FLAG M2 agarose gel (Sigma, A2220) overnight, washed 10 times and eluted with 0.1mg/ml FLAG peptides (ELIM Biopharmaceutical. Protein complexes were resolved in 10% SDS PAGE.

### Masspectrometry

Protein samples were denatured, reduced, and alkylated before overnight trypsinization at 37° C. Then, samples were concentrated, desalted, dried and resuspended in 0.1% formic acid for mass spectrometry analysis.

Digested peptide mixtures were analyzed by LC-MS/MS on a Thermo Scientific LTQ Orbitrap Elite mass spectrometry system equipped with a Proxeon Easy nLC 1000 ultra high-pressure ‘liquid chromatography and autosampler system. Samples were injected onto a C18 column (25 cm x 75 um I.D. packed with ReproSil Pur C18 AQ 1.9um particles) in 0.1% formic acid and then separated with a one-hour gradient from 5% to 30% ACN in 0.1% formic acid at a flow rate of 300 nl/min. The mass spectrometer collected data in a data-dependent fashion, collecting one full scan in the Orbitrap at 120,000 resolution followed by 20 collision-induced dissociation MS/MS scans in the dual linear ion trap for the 20 most intense peaks from the full scan. Dynamic exclusion was enabled for 30 seconds with a repeat count of 1. Charge state screening was employed to reject analysis of singly charged species or species for which a charge could not be assigned.

Raw mass spectrometry data were analyzed using the MaxQuant software package (version 1.3.0.5)(57). Data were matched to the SwissProt mouse protein sequencess (downloaded from UniProt on 7/19/2016). MaxQuant was configured to generate and search against a reverse sequence database for false discovery rate (FDR) calculations. Variable modifications were allowed for methionine oxidation and protein N-terminus acetylation. A fixed modification was indicated for cysteine carbamidomethylation. Full trypsin specificity was required. The first search was performed with a mass accuracy of +/− 20 parts per million and the main search was performed with a mass accuracy of +/− 6 parts per million. A maximum of 5 modifications were allowed per peptide. A maximum of 2 missed cleavages were allowed. The maximum charge allowed was 7+. Individual peptide mass tolerances were allowed. For MS/MS matching, a mass tolerance of 0.5 Da was allowed and the top 6 peaks per 100 Da were analyzed. MS/MS matching was allowed for higher charge states, water and ammonia loss events. The data were filtered to obtain a peptide, protein, and site-level FDR of 0.01. The minimum peptide length was 7 amino acids. Results were matched between runs with a time window of 2 minutes for technical duplicates. All precursor (MS1) intensities of valid peptide matches were quantified by the Maxquant LFQ algorithm using the match between runs option to minimize missing values. For statistical analysis, the quantitative change of peptides that were uniquely assigned to protein isoforms across all IPs were compared with the MSstats R package (v. 2.3.4)(58). Briefly, all peak intensities were Log_2_-transformed and their distributions were median-centered across all runs using the scale option. All remaining missing intensity values were imputed by setting their value to minimal intensity value per run, as an estimate for the MS Limit Of Quantitation. The normalized dataset was then analyzed by fitting a mixed effects model per protein using the model without interaction terms, unequal feature variance and restricted scope of technical and biological replication. The average change (Log2-Fold-Change) of the model-based abundance estimate was computed by comparing replicates of each differentiation stage against the ES undifferentiated pool. Proteins with a greater than four fold change (Log2 Fold Change > 2) and test p-value < 0.05 were determined as significantly altered during differentiation.

### Nucleosome reconstitution, repositioning and ATPase assay

Recombinant *Xenopus laevis* histone octamers were constituted on a 601-nucleosome positioning DNA(59) as described (60). Nucleosome repositioning and ATP hydrolysis assays were performed as described before(60).

### RNA-Sequencing

Total RNA were isolated from biologically triplicate samples using miRNeasy micro kit with on column DNase I digestion (Qiagen). RNA-seq libraries were prepared with the Ovation RNA-seq system v2 kit (NuGEN). Libraries from the SPIA amplified cDNA were made using the Ultralow DR library kit (NuGEN). RNA-seq libraries were analyzed by Bioanalyzer, quantified by KAPA QPCR and sequenced pair-end 100bp using a HiSeq 2500 instrument (Illumina). RNA reads were aligned with TopHat2(61), counts per gene calculated using featureCounts (62)and edgeR (63) was used for the analysis of differential expression. K-means clustering and pheatmap functions in R were used to cluster and generate heatmaps. GO enrichment analysis were performed using GO_Elite(64).

### ChIP-Seq

Chromatin immunoprecipitation were performed according to O’Geen et al. (65) Briefly, cells at double crosslinked with 2mM disuccinimidyl gulatarate (DSG) and 1% formaldehyde and quenched with 0.125M glycine. Frozen pellets (5×10^7^) were thawed, washed, dounced and MNase digested. Chromatin were sonicated for short time, centrifuged at stored at −80C.

Chromatin (40ug) was diluted to 5 fold, pre-cleared for 2 hrs followed by IP with anti-Brg1 antibody (Abcam, 110641) for 12-16hrs. 5% of samples were used as input DNA. Antibody bound BRG1-DNA complexes were Immunoprecipitated using 25ul of M-280 goat anti-rabbit IgG dyna beads for 2hrs, washed a total of 10 times with increased stringency buffers and eluted eluted with 200ul of elution buffer (10mM Tris.Cl, pH 7.5, 1mM EDTA and 1%SDS). Samples were reverse crosslinked, proteinase k and RNAse A digested and purified using, Ampure beads. To prepare libraries for sequencing, DNA were end repaired, A-tailed, adapter ligated (Illumina TrueSeq) PCR amplified. PCR amplified libraries were size selected and ampure purified. The concentration and size of eluted libraries was measured (Qubit and Bioanalyzer) before sequencing in a NEBNextSeq sequencer.

Reads (single end 75bp) were trimmed using fastq-mcf and aligned to mouse genome mm9 assembly using Bowtie(66). Minimum mapping quality score was set to 30. Statistically enriched bins with a P-value threshold set to 1×10^6^ were determine with input DNA serving as the background model (67). Galaxy (https://usegalaxy.org/) was used to pool multiple replicates to generate browser tracks and tornado plots. GREAT(51) was used to generate gene lists near BRG1 peaks.

### ATAC Sequencing

Assay for Transposase-Accessible Chromatin using sequencing (ATAC-seq) was performed according to Corces et.,al (37) in 2-5 biological replicates. Briefly, 50,000 cells (> 95% viability) were lysed, washed and tagmented for 45 mins and 3hr for CP and CM cells respectively. DNA was purified, and amplified using universal Ad1 and barcoded reverse primers (37). Libraries were purified, quantified and analyzed on bioanalyser and sequenced on a NEB NextSeq 550 sequencer using Illumina NextSeq 500/550 High Output v2 kit (150 cycles). Sequencing image files were de-multiplexed and fastq generated. Reads (paird-end 75bp) were trimmed and aligned to mouse genome mm9 assembly using Bowtie (66) with a minimum mapping quality score of 30. Statistically enriched bins with a P-value threshold set to 1×10^6^ were determined (67). UCSC genome browser and IGV were used to view the browser tracks. Galaxy (https://usegalaxy.org/) was used to pool multiple replicates to generate browser tracks and tornado plots. GREAT was used to generate gene lists near ATACseq sites.

**Fig. S1.**
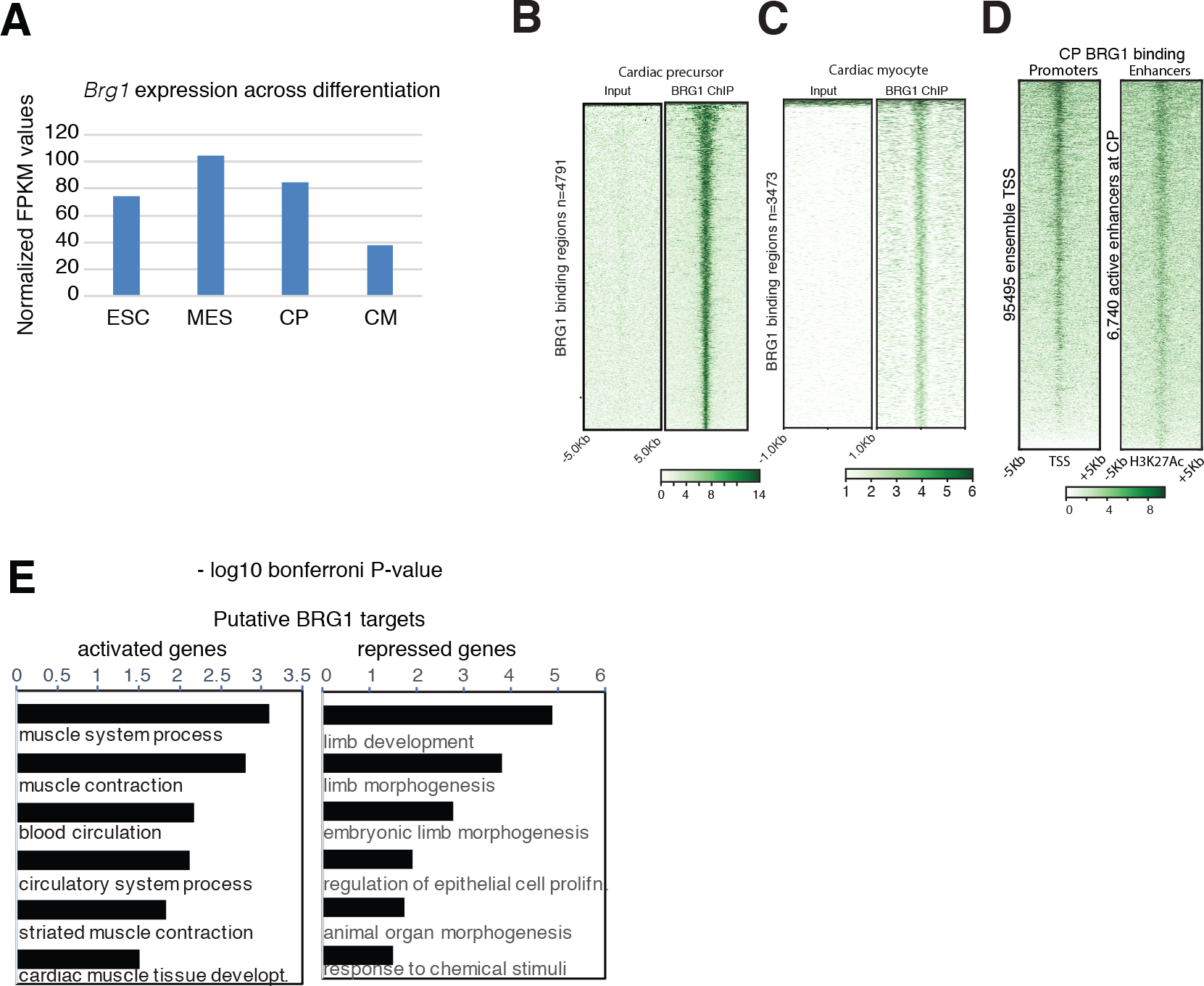
*Brg1* expression and BRG1 genomic binding decreases from CP to CM. (*A*) Expression level of *Brg1* mRNA across indicated four different stages of cardiac differentiation. (*B*) Input and BRG1 ChIP signal over 4791 cardiac precursor BRG1 binding sites. (*C*) Input and BRG1 ChIP signal over 3473 cardiac myocyte BRG1 binding sites. (*D*) BRG1 ChIP signal over TSS of all 95495 emsemble genes and 6740 enahncers in cardiac precursors that are enriched for H3K27ac marks. (*E*) GREAT analysis of biological processes enriched in two flanking genes (within 1Mb) of a BRG1 binding regions at cardiac precursors.

**Fig. S2.**
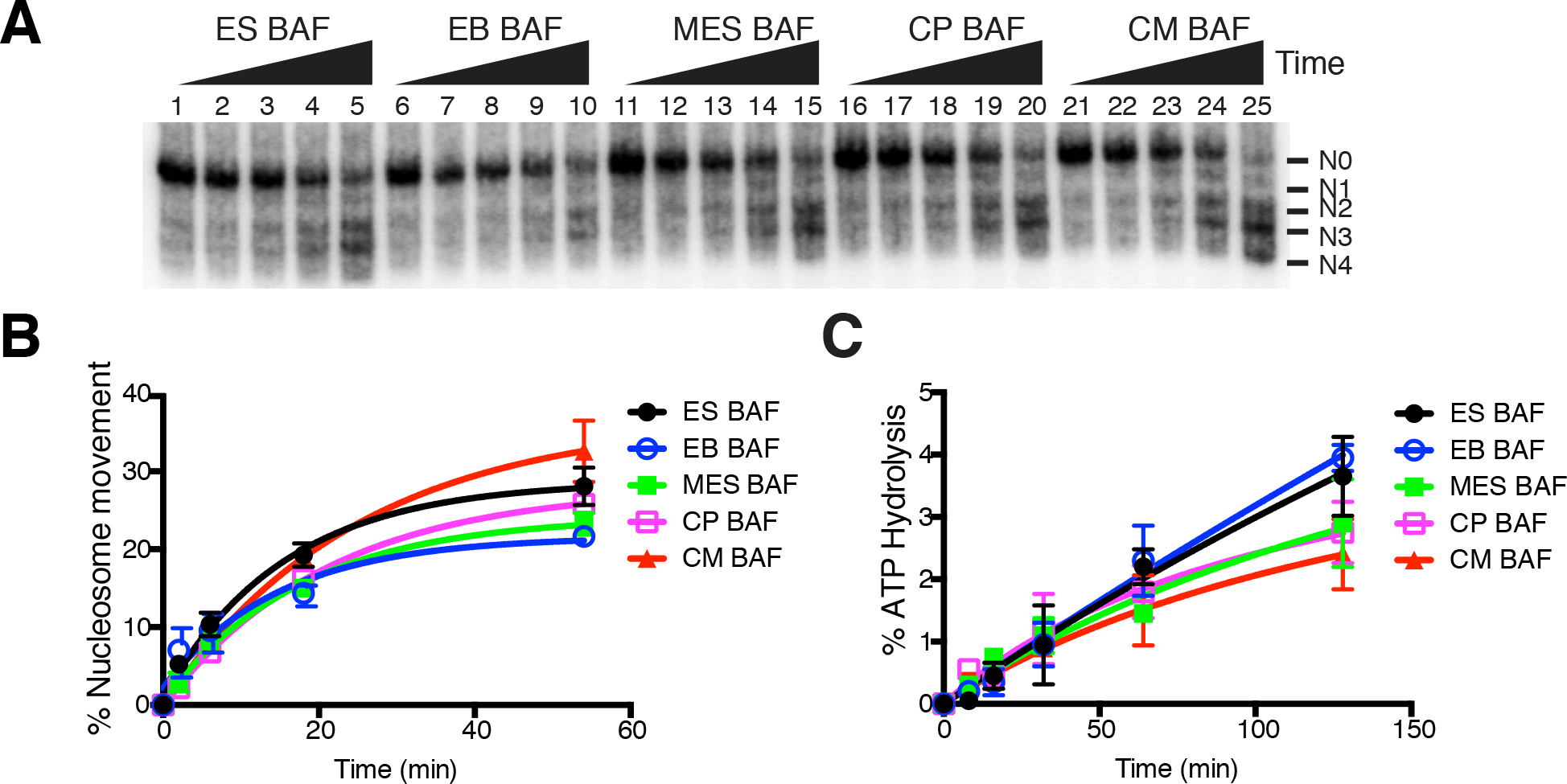
BRG1 containing complexes from different stages of cardiac differentiation do not significantly change nucleosome repositioning or ATPase activities. (*A*) Nucleosome repositioning by BAF complexes isolated at indicated stages of cardiac differentiation. N0 represents the starting nucleosome position. N1-N4 represent BAF displaced repositioned nucleosomes over time. (*B*) Quantification of nucleosome repositioning. Error bars represent s.e.m of two independent replicates. (*C*) Nucleosome stimulated ATP hydrolysis by BAF complexes isolated at different stages of cardiac differentiation. Error bars represent s.e.m of three independent replicates

**Fig. S3.**
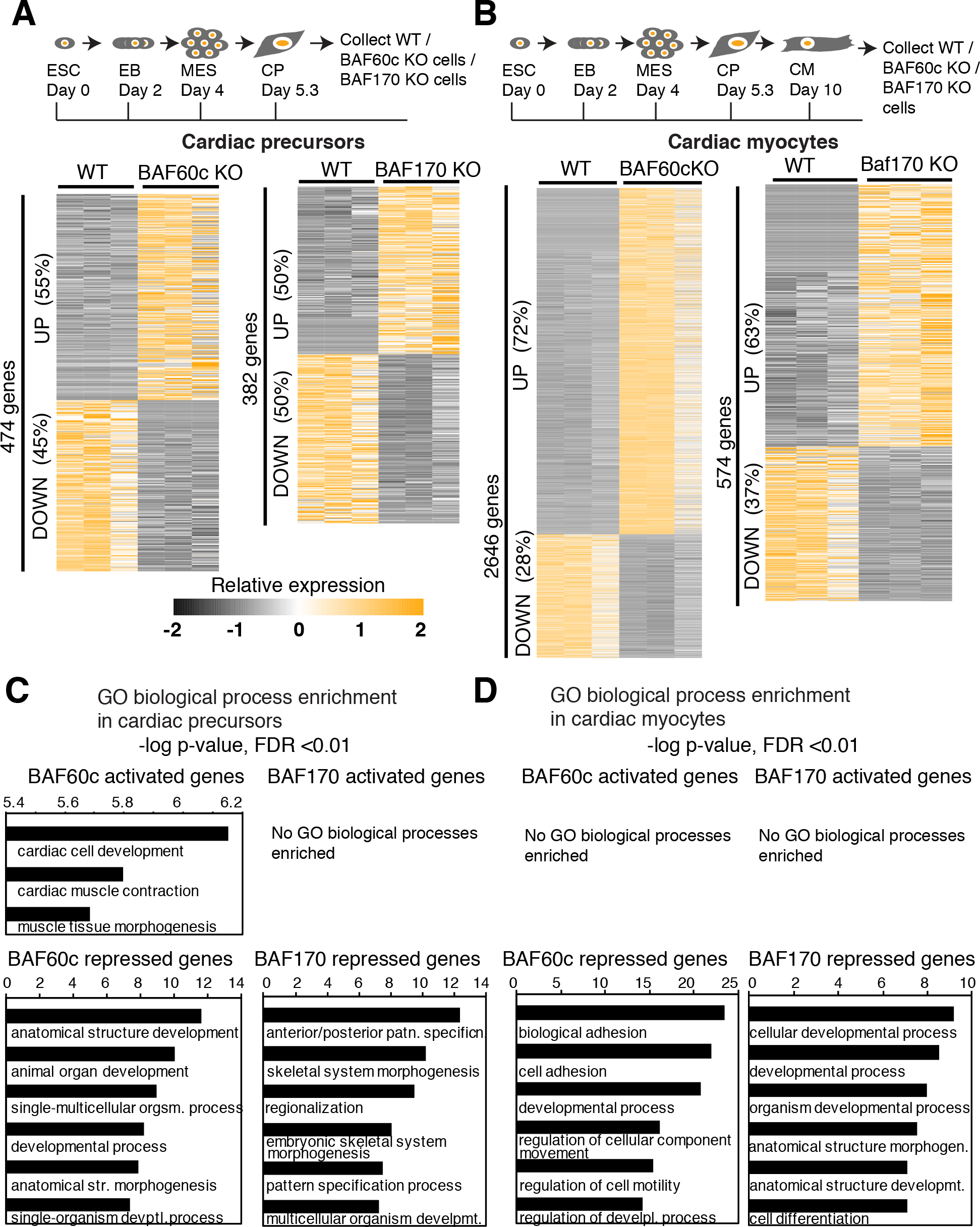
RNAseq analysis of BAF60c KO and BAF170 KO in CP and CM stages of cardiac differentiation. (*A*) Schematics of cardiac differentiation and time point of cell collection for RNAseq. Heat map showing genes up or down regulated in absence of BAF60c (left panel) or BAF170 (right panel) in cardiac precursors. (*B*) Same as A but in cardiomyocytes. GO biological processes enriched in *Baf60c* or *Baf170* KO cells in cardiac precursors (*C*) or cardiac myocytes (*D*) are shown. Processes down regulated in mutants were shown on the top panels and processes upregulated shown on bottom panels.

**Fig. S4.**
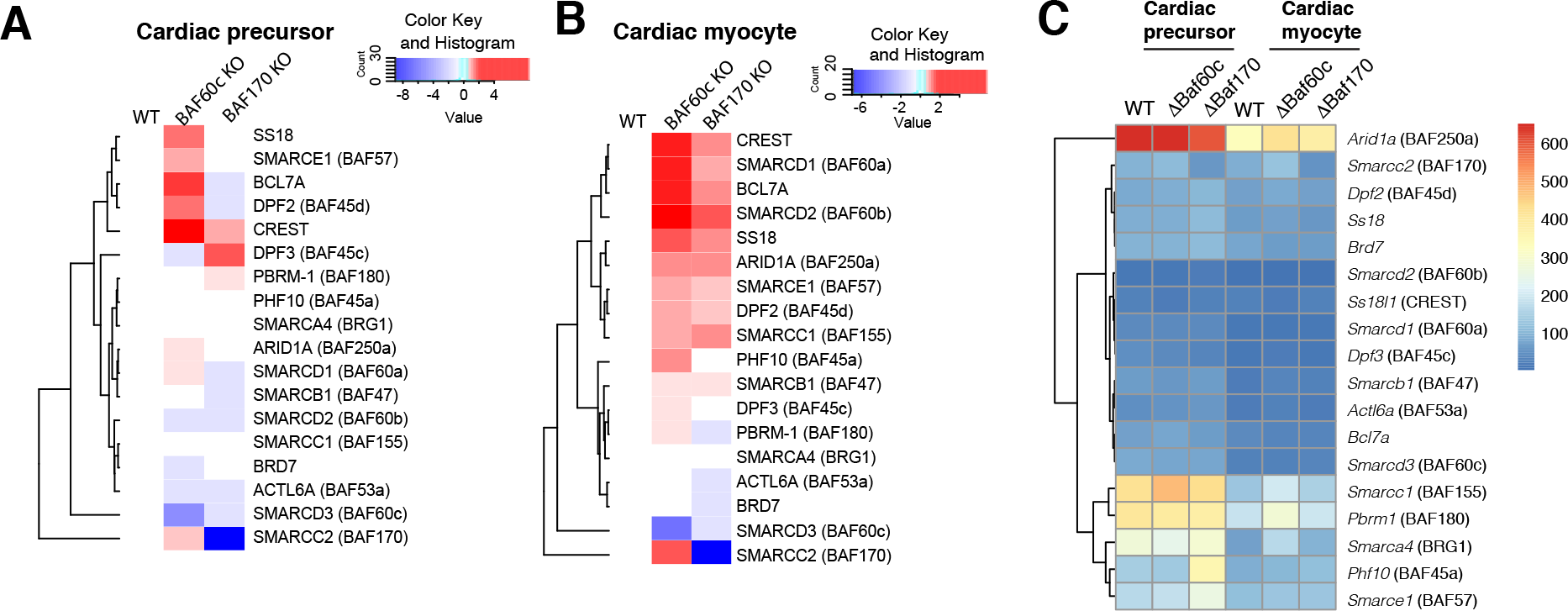
BRG1 complexes form sub-modules in absence of BAF60c and BAF170 Peptide intensities of BRG1 complexes from cardiac precursors lacking BAF60c or BAF170 are normalized to the protein levels of BRG1 and to their WT counterparts. (*B*)Same as *A*, except in cardiomyocytes. Color bar indicates relative association of proteins with BRG1 with blue, white and red representing depletion (less than 1.25-fold), no change (within 1.25-fold) and enrichment (more than 1.25-fold change) in protein abundance respectively. (*C*) Gene expression analysis of BRG1 associated subunits at the indicated stages in absence of BAF60c or BAF170 were plotted for the indicated genotypes at CP and CM. Color bar shows the normalized median FPKM expression values from three biological replicates.

**Supplementary Table S1:**
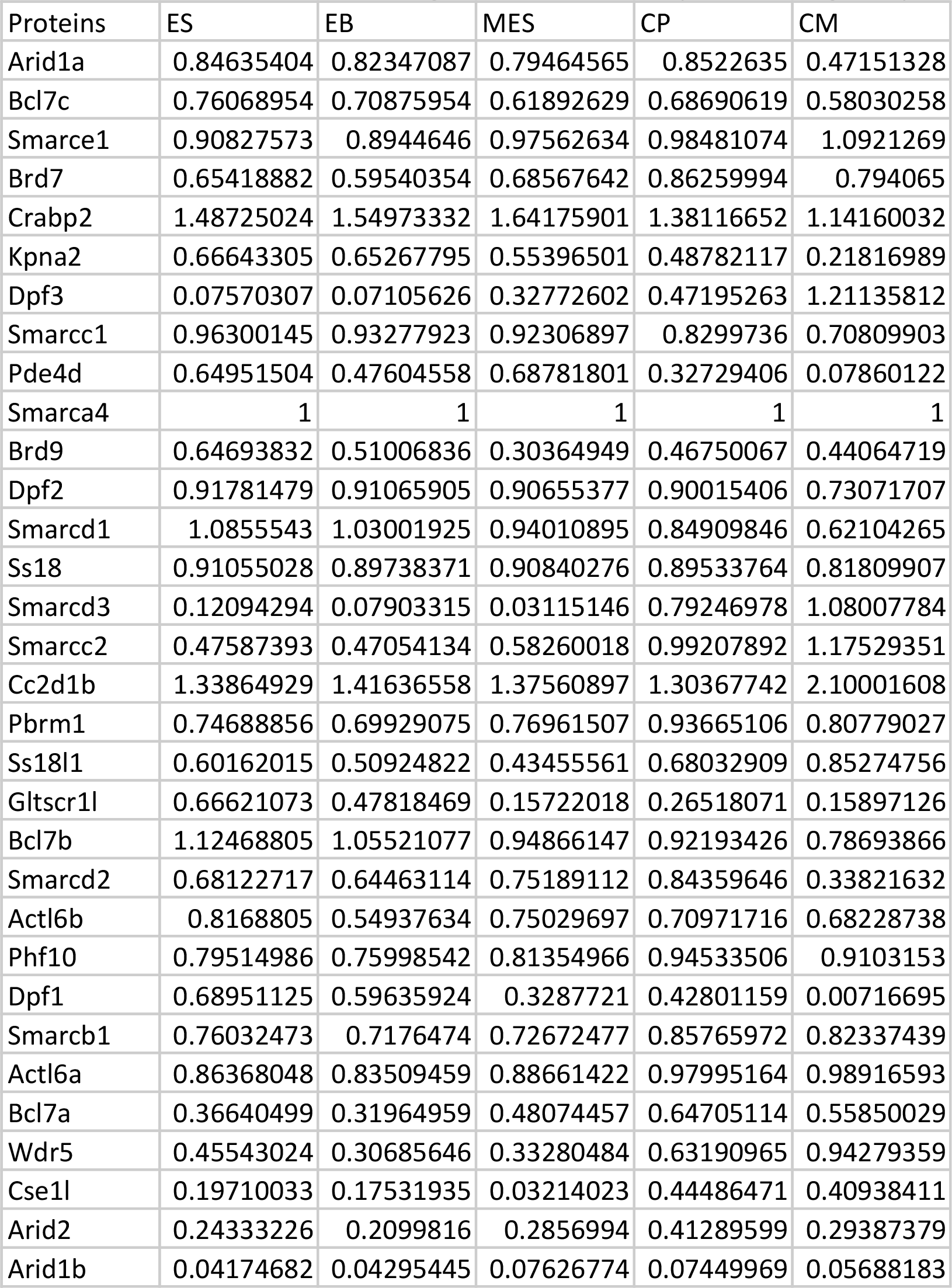
Protein intensities (p<0.05, >4fold over mock control) normalized to BRG1 protein intensities at each stages of differentiation (related to Figure 1)

